# Interactive Analytics for Very Large Scale Genomic Data

**DOI:** 10.1101/035295

**Authors:** Cuiping Pan, Nicole Deflaux, Gregory McInnes, Michael Snyder, Jonathan Bingham, Somalee Datta, Philip Tsao

## Abstract

Large scale genomic sequencing is now widely used to decipher questions in diverse realms such as biological function, human diseases, evolution, ecosystems, and agriculture. With the quantity and diversity these data harbor, a robust and scalable data handling and analysis solution is desired. Here we present interactive analytics using public cloud infrastructure and distributed computing database Dremel and developed according to the standards of Global Alliance for Genomics and Health, to perform information compression, comprehensive quality controls, and biological information retrieval in large volumes of genomic data. We demonstrate that such computing paradigms can provide orders of magnitude faster turnaround for common analyses, transforming long-running batch jobs submitted via a Linux shell into questions that can be asked from a web browser in seconds.

## Main Text

Genomic sequencing projects have gone from sequencing a few genomes to hundreds or thousands of genomes in the past decade ^1–11^. The massive amount of data generated from these studies has presented challenges ranging from affordable long-term storage, controlled data sharing, flexible data retrieval, fast and scalable data processing, and interactive downstream mining of biological information. Recently, jointly analyzing genomic data from multiple studies ^12^ and with other types of data ^13^ has proven to be invaluable in improving study power and thus yielded important new discoveries. These data integration efforts have highlighted the need for a scalable analysis platform that can combine various information sources. Standard public tools, run on local files on fixed-size computer clusters, do not readily scale for large studies. Public cloud platforms can provide part of the solution, with sufficient computing capacity for large batch analysis jobs, and have demonstrated early potential to grow into mature solutions for handling large-scale biology and health data ^14–17^. Another part of the solution may be new interactive analytics frameworks based on distributed computing approaches, such as Dremel ^18, 19^, a query engine based on a columnar database and available in both open source and commercial implementations. Here we present cloud platform-based data processing and Dremel-based interactive analytic solutions for very large scale genomic data. We have built such solutions according to the standards of Global Alliance for Genomics and Health, aiming to help achieve the goal of creating a common framework for securely sharing and analyzing genomic and clinical data.

The dataset of this study contains 460 whole human genomes sequenced with 101 base-pair paired-end reads on the Illumina HiSeq 2000 with an average genome coverage of 50x, totaling 48 terabytes of pre-aligned reads presented in BAM format. We processed these data using our automated BWA and GATK Unified Genotyper pipeline ^20^ as implemented on an in-house computer cluster at Stanford University (Figure 1 step i [Figure 1.i]). Each genome was processed individually, and we turned on the option in GATK to detect both reference and variant bases, yielding in total 9 terabytes of genotypic information presented in compressed VCF format (Methods). To enable cloud-based analysis, we uploaded these VCF files to a storage bucket (Figure 1.ii). As the calls were presented for each position individually and large amount of them were reference calls, we transformed the calls for every genome from VCF to gVCF format in which variant calls were retained in single entries and consecutive reference calls were collapsed into single intervals (Methods, Figure 1.iii). This transformation reduced data size by 150 times as compared to their uncompressed VCF formats, therefore largely reduced storage and facilitated retrieval of variant information. During the transformation we also tagged the calls by quality, marking the low quality reference and variant calls for downstream processing. Subsequently, we imported the gVCF files into a single variant set called genome_calls, which followed the standard of variant set in Global Alliance for Genomics and Health. Then we exported genome_calls to an interactive analytics system based on Dremel for further analysis (Figure 1.v).

**Figure 1:**
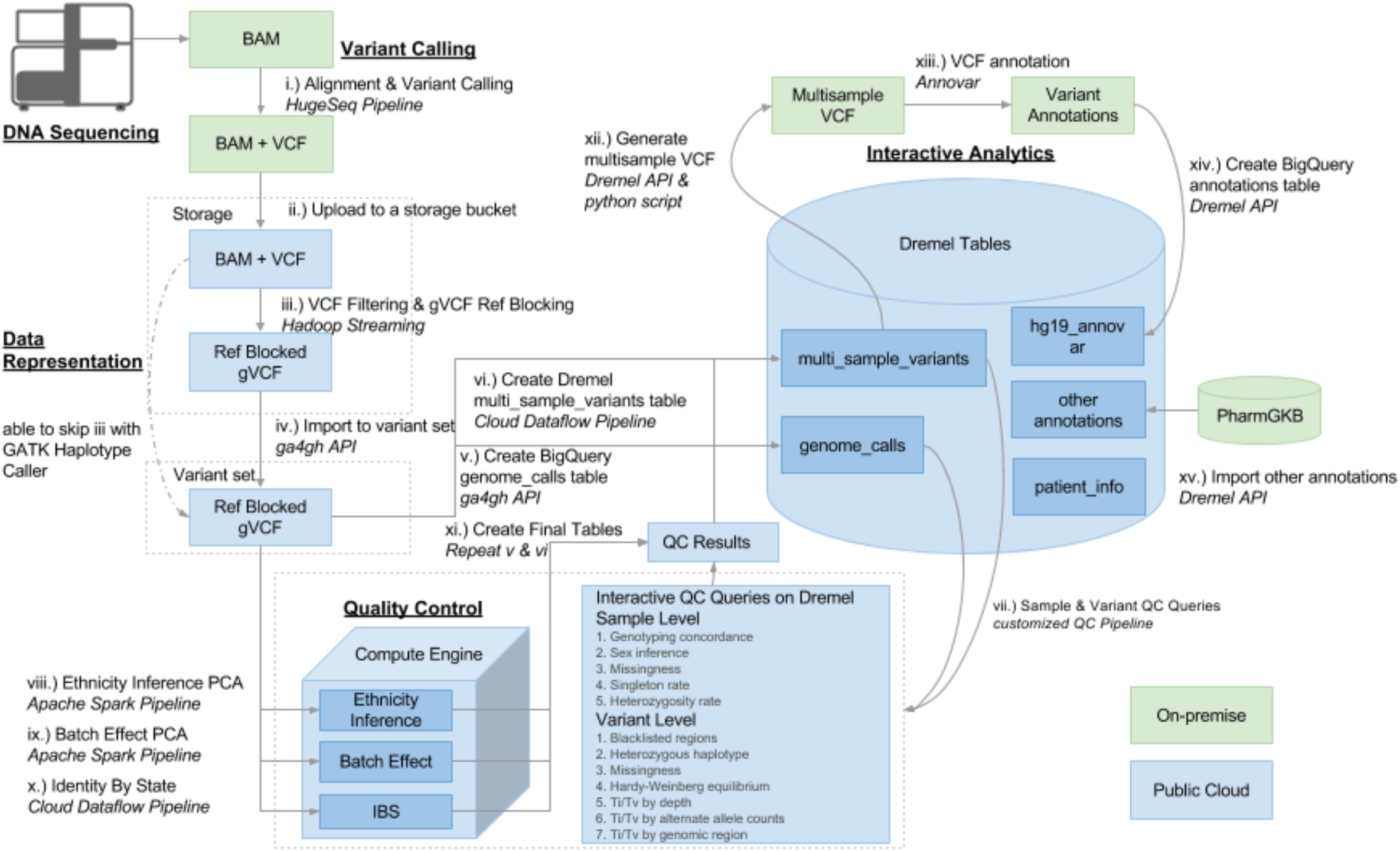
Bioinformatics workflow describing how we utilized on-premise computer cluster for preparing genomic input files (in green) and public cloud infrastructure for carrying out file format transformation and quality controls (in blue). The derived data were imported into Dremel for interactive analysis. i.) BAM files provided by Illumina Sequencing Services were reanalyzed using the HugeSeq pipeline to perform primary genome analysis. Hugeseq comprises BWA and GATK UnifiedGenotyper among other tools. ii.) VCFs output from step i. were uploaded to a cloud storage bucket. iii.) The VCFs were then processed with a Hadoop Streaming job in order to group reference calls into reference-matching blocks and flaq low quality loci. iv.) gVCFs with reference matching blocks were imported into a Global Alliance for Genomics and Health (ga4gh) variant set. v.) A Dremel table resembling a gVCF was generated by exporting the contents of the variant set using the ga4gh API. vi.) A Dremel table resembling a multisample VCF was generated using a Cloud Dataflow pipeline that read data from the variant set and output a Dremel table. vii.) Sample and variant level quality control queries were run through the Dremel API using a python pipeline. viii.) Ethnicity was inferred for the genomes in this study by running a Apache Spark pipeline on 100,000 randomly selected sites from dbSNP for the 460 genomes in this study combined with the 1092 genomes from 1,000 Genomes phase 1^10^. The pipeline reads data from the variant set through the ga4gh API and outputs the PCA results to a storage bucket. ix.) The same Apache Spark pipeline from the previous step was run on all sites of the 460 genomes from this study to test for sequencing batch effects. x.) Identity by State (IBS) was run for all genomes comparing each genome to all other genomes to identify unexpected relatedness. xi.) Final genome_calls and multi_sample_variants tables were generated after combining the results from each of the QC steps (vii-x). Low quality loci are flagged in the final tables with a genotype of -1/-1 and low quality samples are removed from the dataset entirely. xii.) A multisample VCF was generated from the multi_sample_variants table by exporting the table and running a python script for formatting. xiii.) Annovar was run on a local HPC to perform variant annotation on all high quality loci. xiv.) A Dremel annotations table was created from the Annovar results. xv.) Other genomic databases such as PharmGKB were imported as Dremel tables to perform additional annotation.

Dremel was originally built by Google to analyze petabyte scale log files. Available implementations include open source databases such as Apache Drill and Cloudera Impala, and Google BigQuery. We adapted a Dremel schema for scalable variant mining to achieve interactive performance for genomic queries. The schema we designed organizes variant calls in a tree-structured schema analogous to the VCF format but with nested and repeated fields (Figure 2). It lists definitive parameters such as chromosome names, positions, and reference bases as single variables, and diversified parameters such as alternate bases as repeated variables to accommodate multiple scenarios that appeared in the cohort. We also hierarchically organized records into flat and nested fields. For example, *call* is nested to record multiple parameters from read alignment and variant calling, as distinct from other fields such as chromosome name and dbSNP ID. This hierarchical structure facilitates queries for cohort information, e.g. counting total unique variants in the cohort, by traversing less data and therefore reducing both computation time and cost. On other occasions where we need to examine the calls from multiple aspects such as depth, mapping quality score, and variant score, we can unfold the nested fields to access details. Due to these repeated and nested fields, parameters of read alignment and variant calling as recorded in the original VCF files can be maintained for each genome and accessed in a single table. Our schema is designed to be intuitive while enabling interactive response to typical queries in a large scale research project.

**Figure 2.**
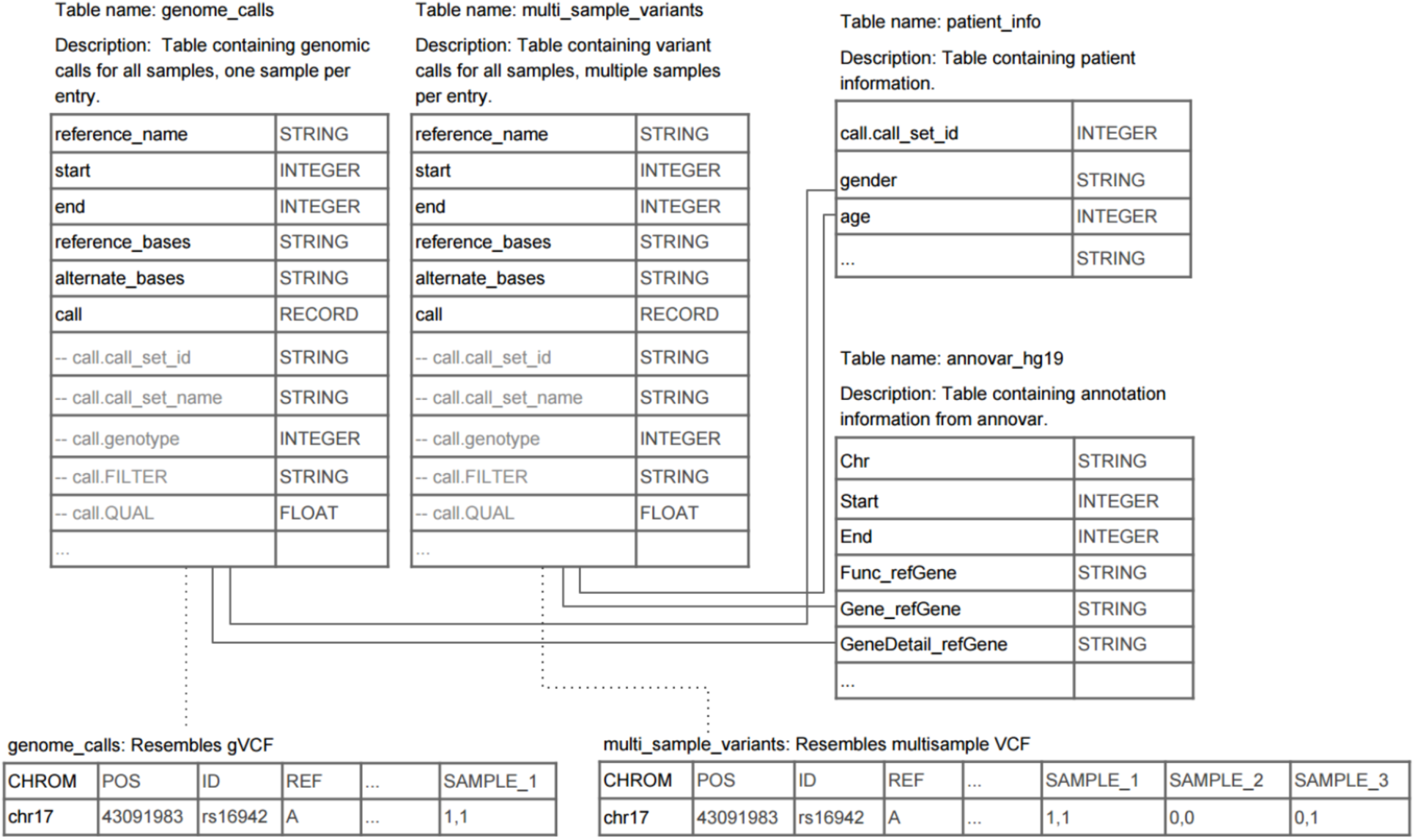
Dremel database schema. Column information for genome_calls, multi_sample_variants, patient_info, and annovar_hg19 tables. Example table data is shown at the bottom for genome_calls and multi_sample_variants.

The table genome_calls contains all reference calls and variant calls. To facilitate variant-centric analysis, we generated the table, multi_sample_variants, based on the genome_calls to document calls for all the variable sites (Figure 1.vi). These are the positions where a variant was detected in any of the genomes in the cohort. Positions where only reference calls were detected were omitted, hence the multi_sample_variants resembles the commonly accepted multi-sample VCF file but with repeated and nested fields.

To derive a list of high quality samples and variants for downstream data mining, we implemented comprehensive quality control (QC) metrics over the SNVs and INDELs using mostly SQL queries. Altogether there are three levels of quality controls: data processing QC, sample health QC, and variant health QC (Figure 3; Supplemental Table 1; Figure 1.iii, vii-x). The data processing QC was implemented during the data transformation step, as mentioned above. In this step, QC of variant calls largely relied on variant quality score recalibration (VQSR) in GATK, which studied the distribution of a variety of parameters for the dbSNP variants identified in our cohort, and expanded the true set to the previously unknown variants ^20^. For reference calls, we hard coded the filtering criteria (Methods). In the tables genome_calls and multi_sample_variants, high quality calls retain their original genotypes (REF: 0/0, VAR: 0/1, 1/1 or 1/2), whereas low quality calls were changed to -1/-1 genotypes (Supplemental Table 2). At the levels of sample health and variant health QC, we considered principles for individual and population genomics. Sample health QC identifies problematic genomes by assessing potential batch effect, unexpected relatedness (e.g. IBS and inbreeding coefficients), concordance to the SNP array data, concordance with phenotypic data (e.g. sex and ethnicity), and distribution of the variants within the cohort (e.g. missingness, singleton, and heterozygosity). Problematic genomes identified from this step were removed entirely from the dataset. The remaining genomes were collected for variant health QC, which aimed to detect problematic variants that 1) mapped to genes that were previously known to have higher failure rate of alignment and variant calling (blacklisted genes ^22^), 2) were mistakenly called as heterozygous in sex chromosomes in male genomes (heterozygous haplotype), 3) were undetected in a significant portion of the cohort (missingness), or 4) failed population genetics tests (HWE and Ti/Tv). Variants failing QC were tagged in the final table. Code for each step is published online (GitHub 1, 2), and the QC process was automated using a custom pipeline that executes queries through a web-based Application Program Interface (API) (GitHub 3).

**Figure 3.**
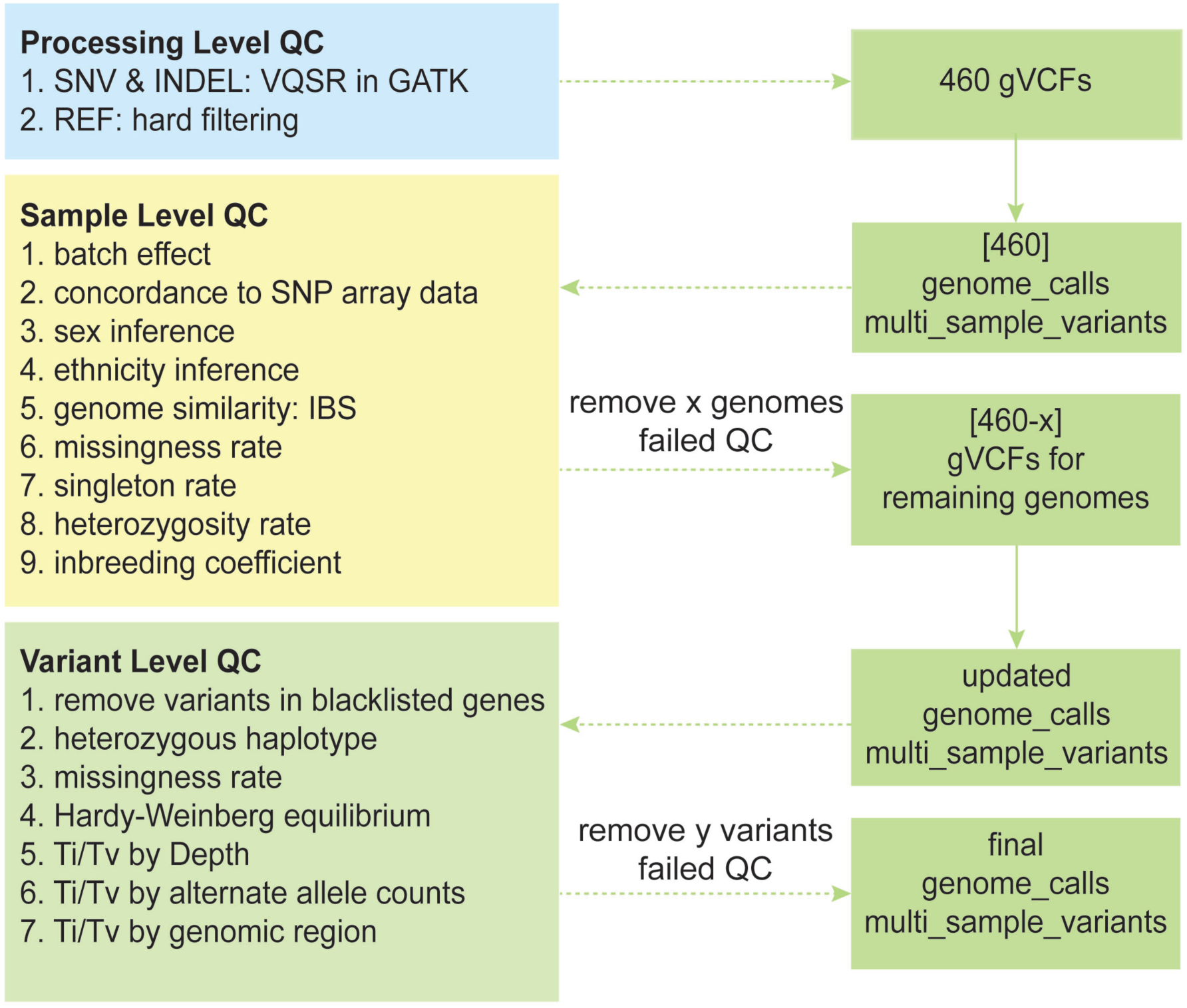
A three-level quality control for large scale genomic sequencing data.

To test the accuracy of our QC steps as implemented in Dremel, we extracted all calls on the BRCA1 gene and compared the QC results with those derived from VCFtools, Plink, and PlinkSeq (Methods: Query validation). Our point-to-point comparison shows very close concordance with those commonly used tools (GitHub 4). We found that most of the few differences were explained by the academic tools disregarding multi-allelic sites. We attempted to better handle multi-allelic sites where appropriate. We then applied these QC steps to our whole genome sequencing cohort (Supplemental Figure 1), and results were summarized in Supplemental Figure 2 and Supplemental Table 3.

After extensive QC on our 460 human genome cohort, we derived in total 446 genomes with 27,076,770 good quality SNVs and 640,341 INDELs. We developed SQL-like, interactive queries to obtain statistical, biological, and medical information. For cohort statistics, we queried these genotypes at various levels to understand how the genotypes distributed within this population (Supplemental Figure 3). We assessed callable rates and a typical scenario in a single genome is displayed in Supplemental Figure 4. We calculated genomic statistics such as Ti/Tv, Het/Hom, novel SNVs and INDELs that were not reported in dbSNP, and private SNVs and INDELs (Supplemental Figure 5). After computing variant allele frequencies within the cohort (Supplemental Figure 6) we observed that close to half of the SNVs were found in only 1–3 genomes, likely due to the fact that these genomes were deeply sequenced. We queried for regions where variation occurred more frequently, the so-called “mutational hotspots”. As an example, we summed the number of SNVs for every 1 kb for P53 gene in each genome (Supplemental Figure 7). We also implemented more comprehensive analytics such as genome wide association studies to facilitate discovery of trait-associated SNVs (GitHub 5). To study the potential impacts the variants could have on biological functions, we annotated the variants using Annovar and uploaded the results to Dremel (Methods, Figure 1.xiii-ix). By querying the annotated features, we identified variants in different genic regions and when normalized by chromosome lengths, we observed that chromosome 19 tended to have denser variation related to coding events locating on exonic, splicing, UTR5 and UTR3 regions (Supplemental Figure 8). There have been increasing efforts to understand medical implications from genomic sequencing data ^23–26^. We imported into Dremel the pharmGKB database ^27^ and queried for all variants in our study that could affect Warfarin dosage effect (Fig 1.xv). Results show that a third of our study subjects should be given a lower dose of Warfarin (Supplemental Figure 9).

Dremel, like other database systems, allows a researcher to join their data with other public and private sources. In our experimentation, we uploaded the calls of the 1000 Genomes Phase 1 project and compared allele frequency distributions in our cohort and in the European populations of the 1000 Genomes Project (Supplemental Figure 10). We observed that in the lower allele frequency categories, our whole genome sequencing study contributed significantly more variants. This can be expected as our deep sequencing was able to identify more rare variants.

Scalability and cost are two major concerns for many computational platforms when hundreds to tens of thousands of genomes need to be analyzed. Our tests on 5 to 460 genomes in Dremel are shown in Figure 4. The underlying distributed storage and elastic computer cluster is built for scalability, and we designed the schema to specifically address interactive variant mining for large scale genomic studies, with the expectation that the platform will scale up well to orders of magnitude more genomes. Runtime comparison between Dremel and other public tools are provided and we observed that as we went from 5 to 460 genomes (90x), the run-time continued to be interactive (i.e. in seconds) whereas the public tools run-time increased from interactive mode (i.e. 1–2 minutes) to batch mode (i.e. 1–2 hours) (Methods, Supplemental Table 4). While costs of platforms are hard to compare, we found Dremel to be marginally more expensive when compared to public tools running on fixed-size computer clusters running near capacity (Methods, Supplemental Table 5). However, any hypothetical cost savings must be traded off against the orders of magnitude longer total wall time and the inability to do interactive queries for data exploration.

**Figure 4.**
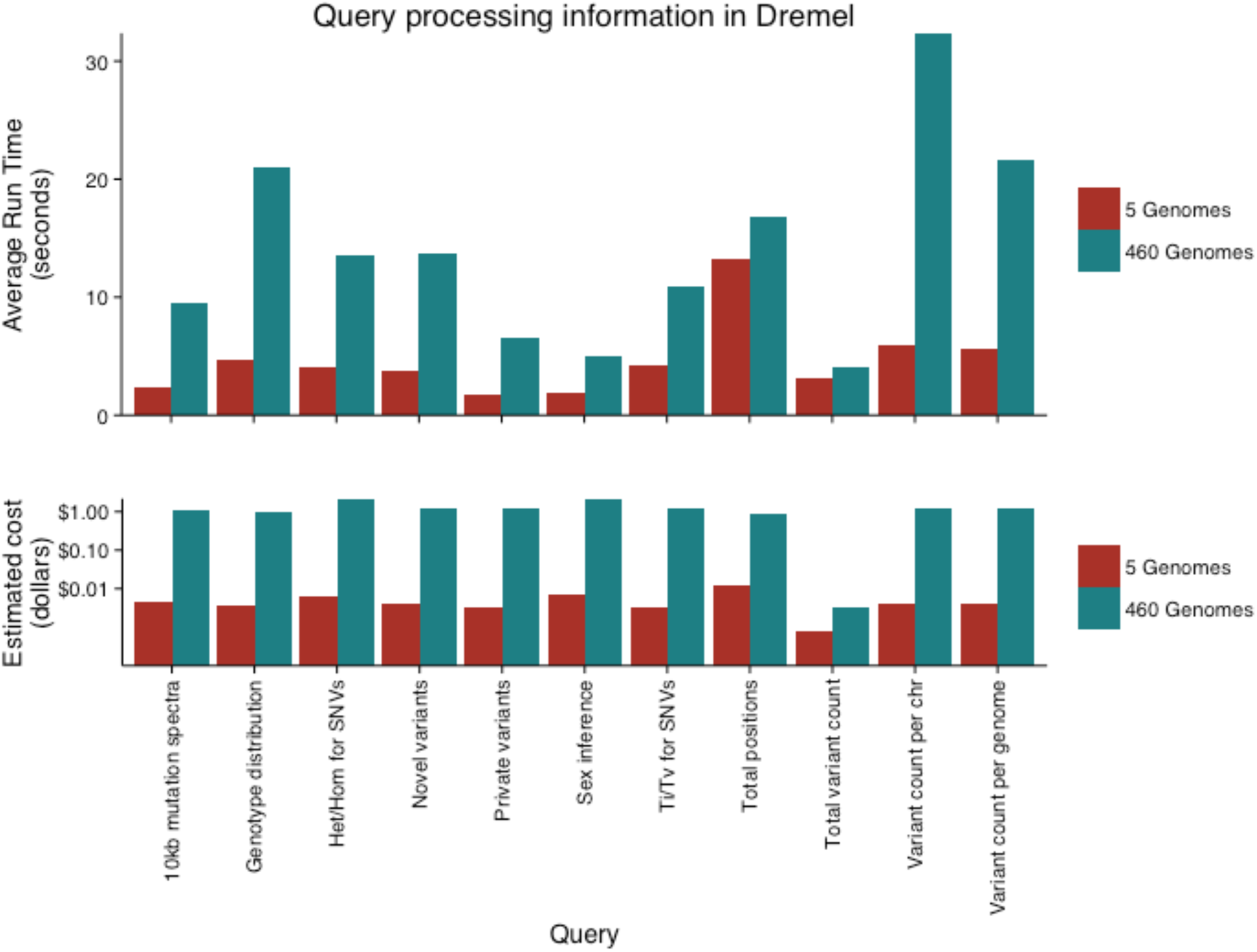
Performance of Dremel in analyzing 5 versus 460 human genomes in this study. Top: Average run times for each query. Bottom: Estimated cost, based on $5/TB of data traversed.

In this study we have presented the computational paradigm combining cloud-based systems and distributed query engines to address the scalability and interactivity in large genomic data analytics. We have demonstrated that such solutions can greatly shorten the cycle of data analysis and hypothesis testing, transforming long-running batch jobs into questions that can be asked from a web browser in seconds. We have also demonstrated, by the examples of calculating IBD and PCA, that the Global Alliance for Genomics and Health API can be used to perform real computation in a future where data is hosted in the cloud and securely accessible with interoperable standards. To our knowledge this is the first implementations of the Global Alliance for Genomics and Health API. We envision that with such APIs, the data processing and exploration may be simplified even further. As an example, the QC Pipeline we built for this study has greatly facilitated the analysis workflow. Even today, tools like Dremel provide flexibility for researchers to build applications beyond what we have demonstrated so far, and tailor to their own research. Though we tested a cloud-based service implementing Dremel, researchers restricted from using public clouds can choose to install and operate one of the multiple implementations on premises, and benefit from much of the performance, subject to the constraints of cluster size and utilization. The performance and scalability of these query engines make them specifically applicable to large sequencing projects.

## Acknowledgement

This work was supported by research grants from the National Institutes of Health (1P50HL083800 and 1R01HL101388) and from the Veterans Affairs Office of Research and Development. We acknowledge the Genetics Bioinformatics Service Center (GBSC), a Stanford School of Medicine Service Center that provides computational infrastructure for genomics research. GBSC provided the dbGaP compliant on-premise cluster and cloud gateway for this research. We acknowledge Bina Technologies Inc (now Roche) for providing technical support for calling genomic variants on Bina servers. We thank Alicia Deng and Hassan Chaib from Stanford University for preparing the DNA for sequencing, Denis Salins and Isaac Liao from Stanford University for bioinformatics support, Elmer Garduno, Danyao Wang, and Samuel Gross from Google for bioinformatics support on Google Cloud, Shannon Rego from Stanford University for discussing results, and David Glazer from Google for insightful advice along the project.

GitHub links:
1. https://github.com/StanfordBioinformatics/mvp_aaa_codelabs/blob/master/Sample-Level-QC.md
2. https://github.com/StanfordBioinformatics/mvp_aaa_codelabs/blob/master/Variant-Level-QC.md
3. https://github.com/StanfordBioinformatics/mvp_aaa_codelabs/blob/master/Comparison.md
4. https://github.com/StanfordBioinformatics/bigquery-genomics-qc
5. https://github.com/googlegenomics/codelabs/blob/master/R/1000Genomes-BRCA1-analysis/sql/gwas-brca1-pattern.sal

